# Population growth poses a significant threat to forest ecosystems: a case study from the Hindukush-Himalayas of Pakistan

**DOI:** 10.1101/2024.04.01.587657

**Authors:** Naveed Alam, Zahid Ullah, Bilal Ahmad, Ahmad Ali, Kashmala Syed

## Abstract

Human population growth and associated increases in anthropogenic activities pose a significant threat to forest ecosystems by diminishing the natural ecosystem services these systems provide. Malam Jabba is located in District Swat Pakistan’s Hindukush-Himalayan temperate zone, which is renowned for ecotourism and skiing and is rich in timber-producing tree species, medicinal plants, and unique biodiversity; however, the majority of Swat Valley’s population relies on Malam Jabba forests for their timber & fuelwood requirements. We examined how the deforestation rate increased with increasing human population density in Pakistan’s Malam Jabba area of the Hindukush-Himalayas. To identify the forest cover, remote sensing, and geographic information systems were used (RS & GIS). The study area’s vegetation was analyzed with the Normalized Divergence Vegetation Index (NDVI) using multitemporal satellite images for the years 1980, 2000, and 2020. The deforestation rate from 1980 to 2020 was then determined using the decay model, and the MATLAB program was used to predict the deforestation rate for the following two decades in relation to the anticipated growth in the human population. Our result revealed that, in the last two decades, the average rate of deforestation increased from 0.7% to 1.93% per year, while the human population of District Swat increased from 1.2 to 2.3 million at a rate of 9% per year. The decay model predicts that the study area’s deforestation rate will increase to 2.5% per year over the next two decades due to the forecasted 11.6% per year population growth rate. Human population growth in District Swat, Pakistan has seriously threatened the nearby forest ecosystems, and a future increase in human population will further accelerate anthropogenic activities like unsustanible tourism, fuel and timber wood collection and urbanization. Based on our results, we recommend that: (i) in addition to reforestation programs and sustainable use of forest resources, the government should implement a long-term forest management plan (ii) where the density of forest cover can be sustained at an equilibrium level dependent of population growth pressure (iii) and areas with extreme human pressure should be designated as most important for in situ conservation approach.

## 1. Introduction

Forests provide ecological and socioeconomic services to sustain approximately 60% of global biodiversity [1] and directly contribute to the fulfillment of human society’s fundamental needs [2,3]. More than two billion people rely on forests for their basic survival needs, including food, fuelwood, shelter, medicine, and forage for domestic animals [4]. Forests play a crucial role in sustaining ecosystem health and ecological services, and more forest cover will be necessary for the current era to mitigate climate changes and environmental challenges [5-7]. Forests are the main source of ecotourism which is the lagest growing industry of the world, which can uplipt the rural economy of the country and provide livelihood apportunities for the local communities [8]. According to the Kyoto Protocol and the United Nations Framework Convention on Climate Change, forests are essential for carbon storage and economic incentives for the country [9-10].

Despite their direct and indirect importance for human well-being, forest ecosystems are diminishing and disappearing at an alarming rate in many parts of the world due to human activities [11,12]. Notably, the natural resources in Hindu Kush-Himalayas are degrading more rapidly than in other parts of the world but still have received less attention internationally than other ecosystems [13,14]. There is a need for routine and timely forest cover assessments to determine current conditions and forecast the effects of future environmental changes [15,16]. Natural resource managers and environmental policymakers have recognized that the conservation of biological diversity depends on protecting and managing intact natural habitats. Such recognition has increased the significance and urgency of international efforts to preserve biodiversity and forest resources [17]. Consequently, the proposed work aimed to detect changes in forest cover over the past four decades to comprehend the pattern of forest cover change, identify the factors responsible for this change, and provide a foundation for forest conservation and management.

Due to the pressing need to quantify forest resources, numerous assessment techniques have been developed in recent years. We outline here a promising workflow that combines the technologies of geographic information systems (GIS) and remote sensing (RS) in order to monitor long-term shifts in the composition of the forest cover [18]. The application of RS necessitates the use of multitemporal and multispectral data to observe the type, quantity, and geospatial location of land use change [19,20], which is essential for studying the effects of anthropogenic activities and other environments variables on plant communities [21]. In contrast, GIS is computer software is used for evaluating, storing, and analyzing RS data [19]. Satellite images are a modern tool for observing ecological disasters and tracking changes in forest cover over a wide area [20]. The ATCOR tool in ERDAS software can correct the topographic and atmospheric errors in satellite images, improve accuracy and reduce error [18]. Normalized Difference Vegetation Index (NDVI) distinguishes green vegetation from other land features based on chlorophyll content [19]. ArcGIS model builder analyzes the NDVI data further for a comprehensive forest cover quantification [20]. For the development of conservation and management policy, large-scale monitoring of forest resources is essential, whereas small-scale monitoring is required for its implementation [22,23].

Human population growth is directly related to economic insecurity, social problems, unsustainable resource consumption, and anthropogenic disturbances in agricultural expansion, unsustanible ecotourism,urbanization, and deforestation [24]. The decay model is the most accurate method for predicting carrying capacity because it considers the strain that growing populations, under linear population growth, places on natural resources, such as an increase in the rate of deforestation and other environmental disturbances [25]. The decay model determines the relationship between exponential population density growth and deforestation.

## 2. Materials and Methods

### 2.1. Study Area

The study area Malam Jabba (34° 47’ 57” N, 72° 34’ 19” E) is situated in Hindukush-Himalayas mountains at elevations between 2000 to 3000 meters above sea level in District Swat of Northern Pakistan. It is a well-known summer resort surrounded by dense, moist temperate coniferous forests containing numerous commercially, medicinally, and ecologically significant plant species, providing a habitat for hundreds of birds and animals. Historically, this region has been home to the Gandhara civilization and its rich culture, where lush green forests and rangelands were likely the primary reasons for their settlements. Malam Jabba forests are experiencing immense anthropogenic pressure in the form of fuel and timber wood collection due to human population growth in the nearby central city of District Swat; heavy grazing, unmanaged ecotourism activities, and agriculture expansion have further threatened the local ecosystem [26,27].

### 2.2. Research Approach and Methodology

Remote Sensing (RS) data were acquired and processed by using Geographic Information System (GIS) to evaluate long-term forest degradation for the years 1980, 2000, and 2020 [18-20]. Data collected by optical satellites with a high resolution are an essential source for various fields of research, including natural resource management, land cover change detection, conservation biology, and population studies [19]. While gathering satellite data, however, factors such as the terrain and atmospheric effects can obscure and otherwise degrade the quality of satellite images [28]. ATCOR tool in ERDAS was used for satellite imagery from 1980, 2000, and 2020 to remove atmospheric and topographic effects. During the atmospheric correction, different values from metadata were utilized, including solar azimuth angle, elevation angle, solar zenith angle, and the angle between the sun and moon [29]. Normalized Divergence Vegetation Index (NDVI) was used to detect forest cover. The NDVI indices were selected from the unsupervised classification menu in ERDAS, and the algorithm was executed on the corrected images from 1980, 2000, and 2020 at regular intervals to obtain NDVI values. Specifically, red and infrared pixel-level brightness values from Landsat images were utilized for this purpose. The results fell from -1 to 1, with positive values indicating more biomass in plants and negative values indicating less biomass. This index was utilized for all vegetation types to obtain forest biomass values. In order to examine NDVI values for forest cover, Global Positioning System (GPS) and Google Earth training samples were gathered. After comparing a training sample with NDVI-classified images from 1980, 2000, and 2020, we obtained maps and NDVI values for the corresponding imageries, with a maximum value of 0.7 for forest classes [30,31].

### 2.3. Data Collection

Primary Data were collected from U.S. Geological Survey (USGS).Landsat-5(TM) images for the year 1980, Landsat-7 (ETM+) images for the year 2000, Landsat-8 (OLI) images for the year 2020, and Digital Elevation Model (DEM) data with a resolution of 30m x 30m from USGS Earth Explorer were collected to evaluate forest degradation for the years 1980, 2000, and 2020 (Table.1) [28]. To confirm satellite data, field trips were conducted to obtain the Global Positioning System (GPS) data on the ground positioning of vegetation cover [30]. Population data was collected from census report-2017 of Pakistan Bureau of Statistics as a secondary data [32].

**Table 1.** Details on the Landsat data with specifications.

| S.no | Platforms | Perio<br>ds | Number of<br>bands | Data size per band<br>(pixels) | Spatial<br>resolution |
| --- | --- | --- | --- | --- | --- |
| 1 | Landsat-5 (TM) | 1980 | 7 | 1334×1511 | 30m |
| 2 | Landsat-7 (ETM+) | 2000 | 8 | 1334×1511 | 30m |
| 3 | Landsat-8 (OLI) | 2020 | 11 | 1334×1511 | 30m |
| 4 | DEM (SRTM) |  | 1 | 1334×1511 | 30m |

### 2.4. Data Processing

A composite was created by stacking the satellite images, each with a unique band structure. Then a projection transformation using the UTM WGS-84 41N coordinate system was applied prior to subsetting. Landsat satellite images were layered, and the “clip” tool in ArcGIS was then used to isolate the study area. The study area was divided into sub-scenes with a pixel size of 1334 x 1511 for each color channel in the original dataset [28]. The NDVI statistics (Table 2) were computed utilizing multitemporal satellite images from 1980, 2000, and 2020. As vegetation’s green color indicates the amount of chlorophyll in leaves, chlorophyll’s spectral response to the electromagnetic spectrum is its reflectance, which is strongest in near-infrared and weakest in the red due to chlorophyll absorption [30].

**Table 2.** Statistics of NDVI images.

| S. No | Year | Min | Max | Mean | St. Dev |
| --- | --- | --- | --- | --- | --- |
| 1 | Oct-1980 | -0.3333 | 1 | 0.5207 | 0.2347 |
| 2 | Oct-2000 | -1 | 0.9813 | 0.4806 | 0.1975 |
| 3 | Oct-2020 | -0.0149 | 0.9981 | 0.5933 | 0.1325 |

ArcGIS’s model builder was used for analyzing NDVI values to calculate forest cover and correct the topographical errors. The tool facilitates the creation, editing, and management of geoprocessing models; it also permits multiple geoprocessing tools to be utilized sequentially in the automation of various tasks. Various tools from the ArcTools box were used to construct the model for this study, including a raster calculator, a clip tool, a raster to polygon tool, a select tool, an added field tool, a geometry calculator, a smoothing tool, a layer to KM tool, and a tool for discovering topological flaws [33,34]. In the study’s model, blue represents the input of satellite images being added with some tool from the toolbox in ArcGIS, and green is the resultant data after calculation in ArcGIS.

### 2.5. Statistical Analysis and Future Forecast

The normalized difference vegetation index for assessing vegetation cover, is represented by the equation [30-31].

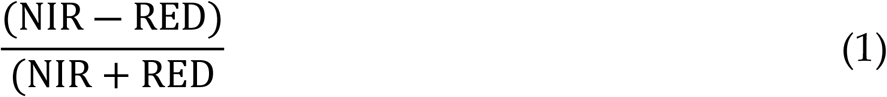

NIR= the brightness value of the near-infrared band.

For predicting future population growth and its effect on forest cover in the form of deforestation, MATLAB 7.10 software was utilized [35]. Using the exponential growth model equation [36,37], we determined the rates of population increase and deforestation for the next two decades.

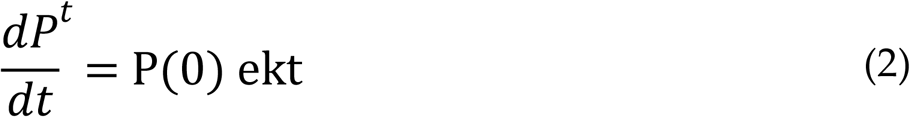

## 3. Results

### 3.1. Forest cover maps

Based on NDVI values, the total forest cover for each year was mapped in 2D and 3D using Arc Map and Arc Scene (Figure 1). In ArcMap, forest maps were generated at the scale of 1: 125,000, while the ArcScene and DEM were used to generate a 3D map of forest cover in the consistent years. The figure shows total forest cover in 8298.039313 polygon area in 1980, 2357.536113 polygon area in 2000, and 609.553980 polygon area in 2020. The polygon area was calculated in ArcGIS, as shown in Table 3 and Figure.2.

**Table 3.** Forest Cover of Study Area & Average Deforestation.

| S. No |  | Forest Cover (ha) | Deforestation (ha) | Average Rate of deforestation |
| --- | --- | --- | --- | --- |
| 1 | Total Forest Cover in October1980 | 15003 | - | - |
| 2 | October 1980 to 2000 | 12859 | 2144 | 0.7%. |
| 3 | October2000 to 2021 | 7894 | 4965 | 1.93% |
| 4 | October 1980 to 2021 | 15003 | 6509 | 1.4% |

**Figure 1.**
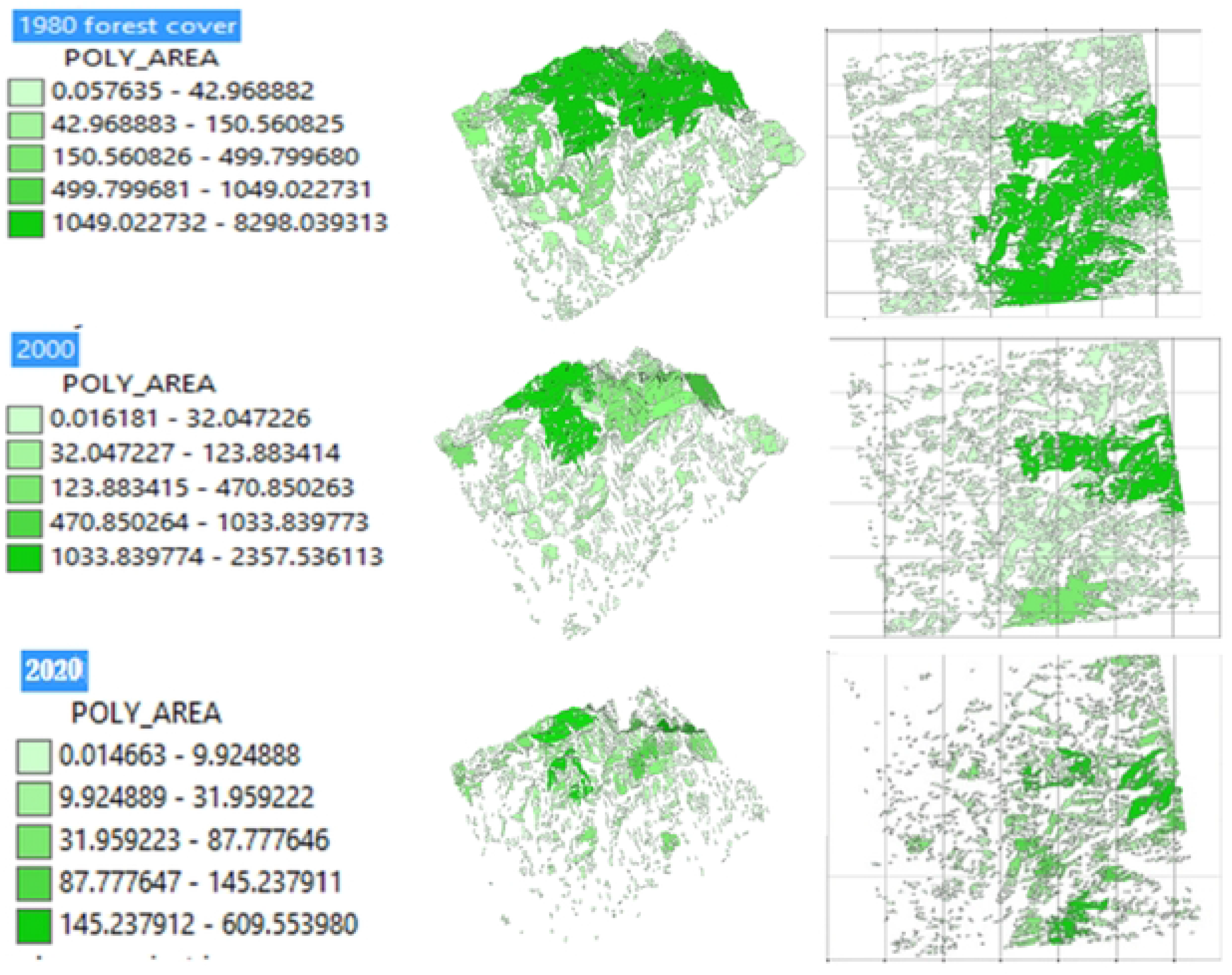
2D and 3D maps showing total forest rover in the years 1980,2000, and 2020.

**Figure 2.**
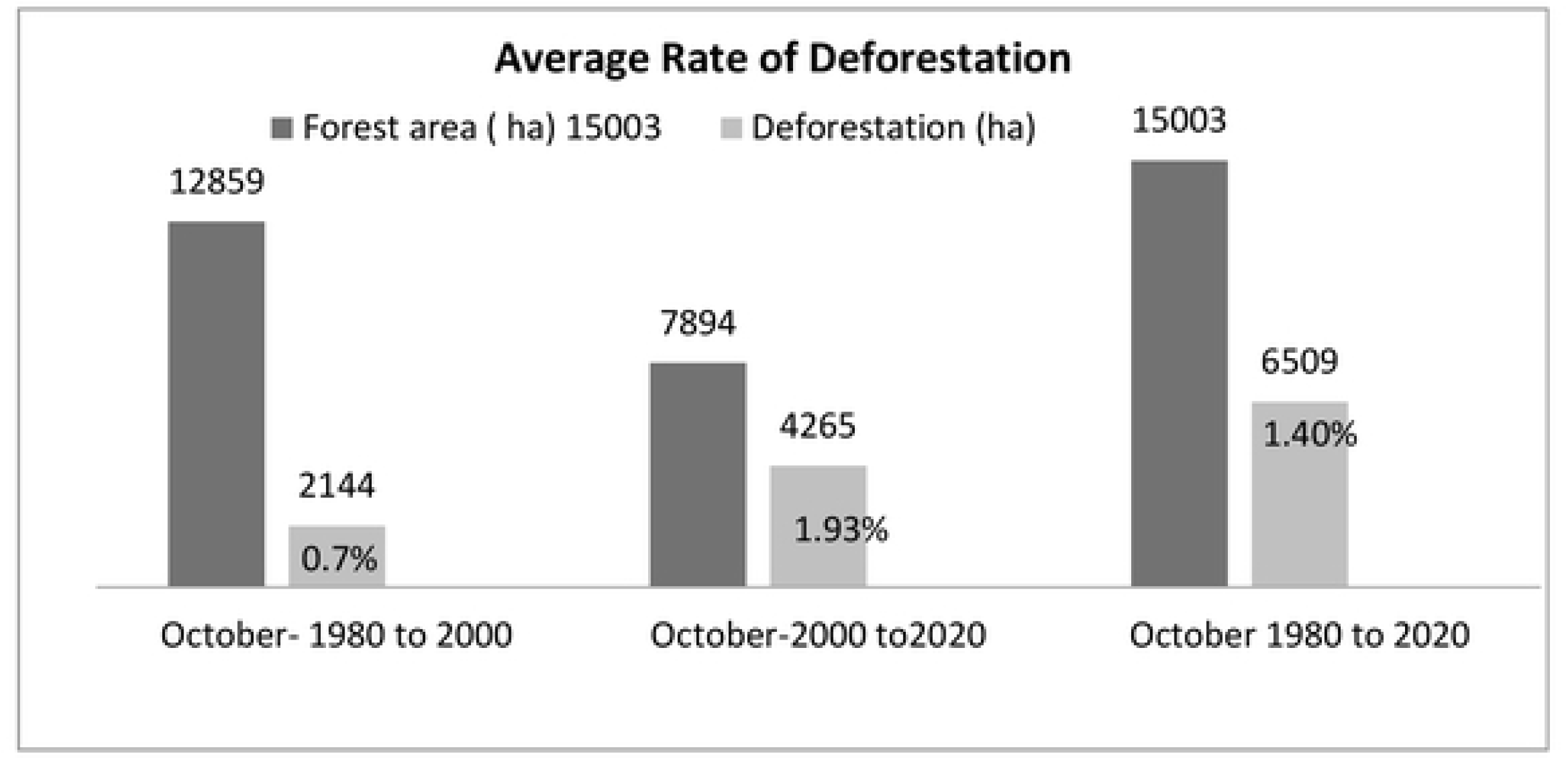
Deforestation rate as measured by total hectares of forest and total forest loss.

### 3.2. Forest Cover Change

Deforestation was defined as the transformation of forest cover into barren land. Our research indicates that between 1980 - 2000, forest cover decreased at an annual rate of 0.7%, from 15 003 ha to a minimum 12 859 ha. From October 2000 to October 2020, the total forest cover decreased at an average annual rate of 1.93 % to reach 7894 ha. From 1980 to October 2020, a total of 6509 ha was affected by deforestation at a rate of 1.40 % annually (Figure 2, Table. 3)

In Southeast Asia, the human population has multiplied in the last two decades, and in the study area it has increased at an average rate of 9.1% (Figure 4). As a result of population growth, pressure on forest resources has increased mainly in the form of timber and fuelwood collection, and our findings show that because of increased anthropogenic activities and unsustainable use of forest resources, forest cover has decreased from 12859 ha to 7894 ha at an average rate of 1.93 ha/year (Figure 2 and 4) The census report-2017 of Pakistan Bureau of Statistics, states that Swat District currently has 2,309,570 residents (Figure 3). The average yearly population growth rate from 1980 to 2000 was 8.9%; this increased to 9.1% for the years leading up to 2020. The exponential/decay model predicts that the population will increase exponentially to 5.7 million by 2040 at 11.6% per year. Anthropogenic activities will rise in tandem with population growth, leading to a deforestation rate increase of 2.5% per year and a loss of forest cover of 4,000 hectares (Figure 6 and 7). In 1851 the total population density was 0.283720 million; in 1961, it was 0.344860 million; in 1972, it was 0.520710 million; in 1981, it reached 0.715940 million; in 1998, it increased to more than a million, and reached to 1.257600 million and finally in 2021 it was recorded 2.309570 million. The decay model shows that District Swat’s population density will reach 5.7 million in 2040. The decay model has also forecasted the forest cover (y-axis) was 15003 hectors in 1980 has been reduced to 12859 hectors in the year 2000, and 7894 hectors in 2020 and will exponentially increase in deforestation will reduce the forest cover to 4000 hectors in 2040 (Figure 6).

**Figure 3.**
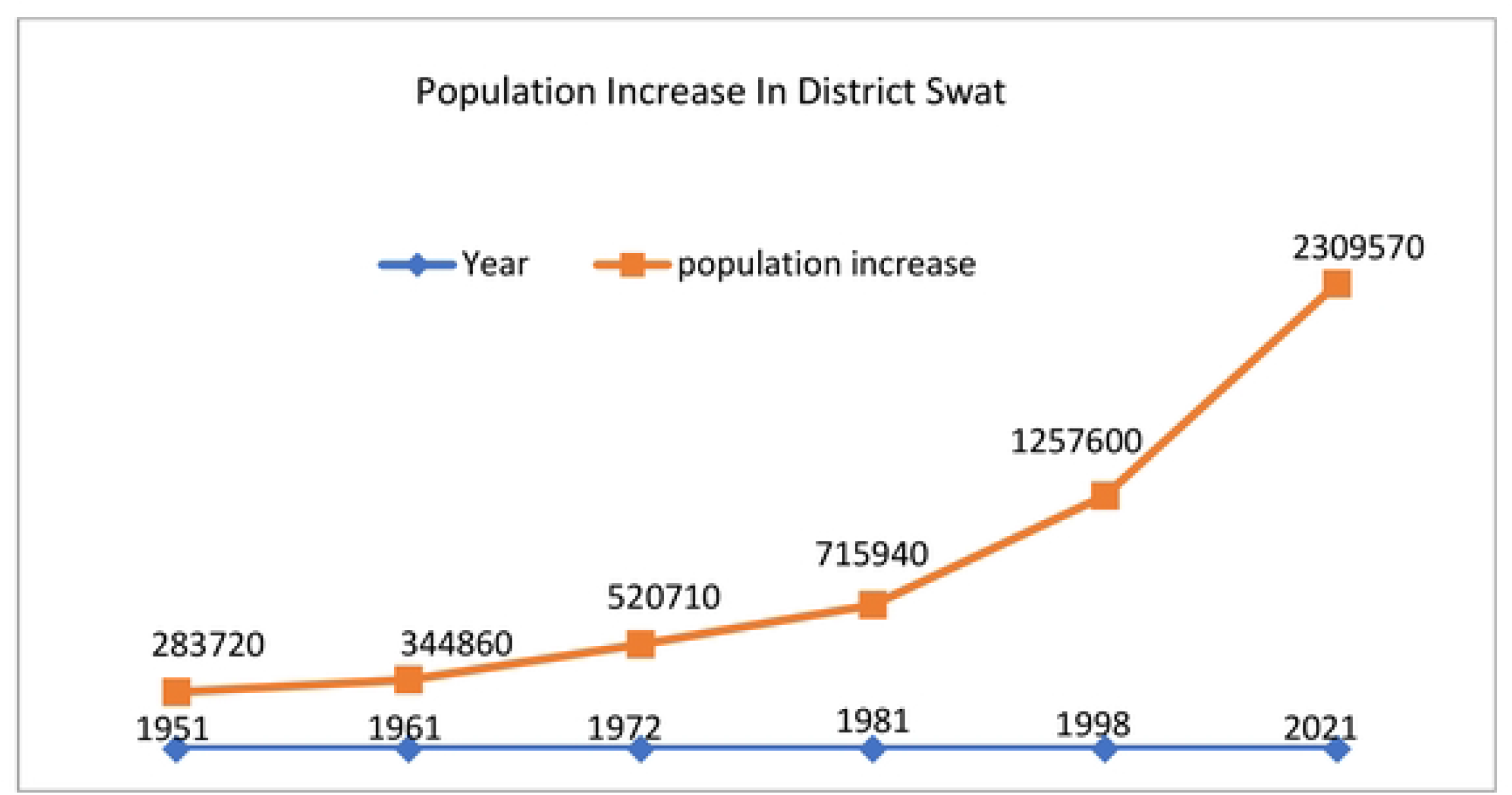
Population Growth within the Study Area.

**Figure 4.**
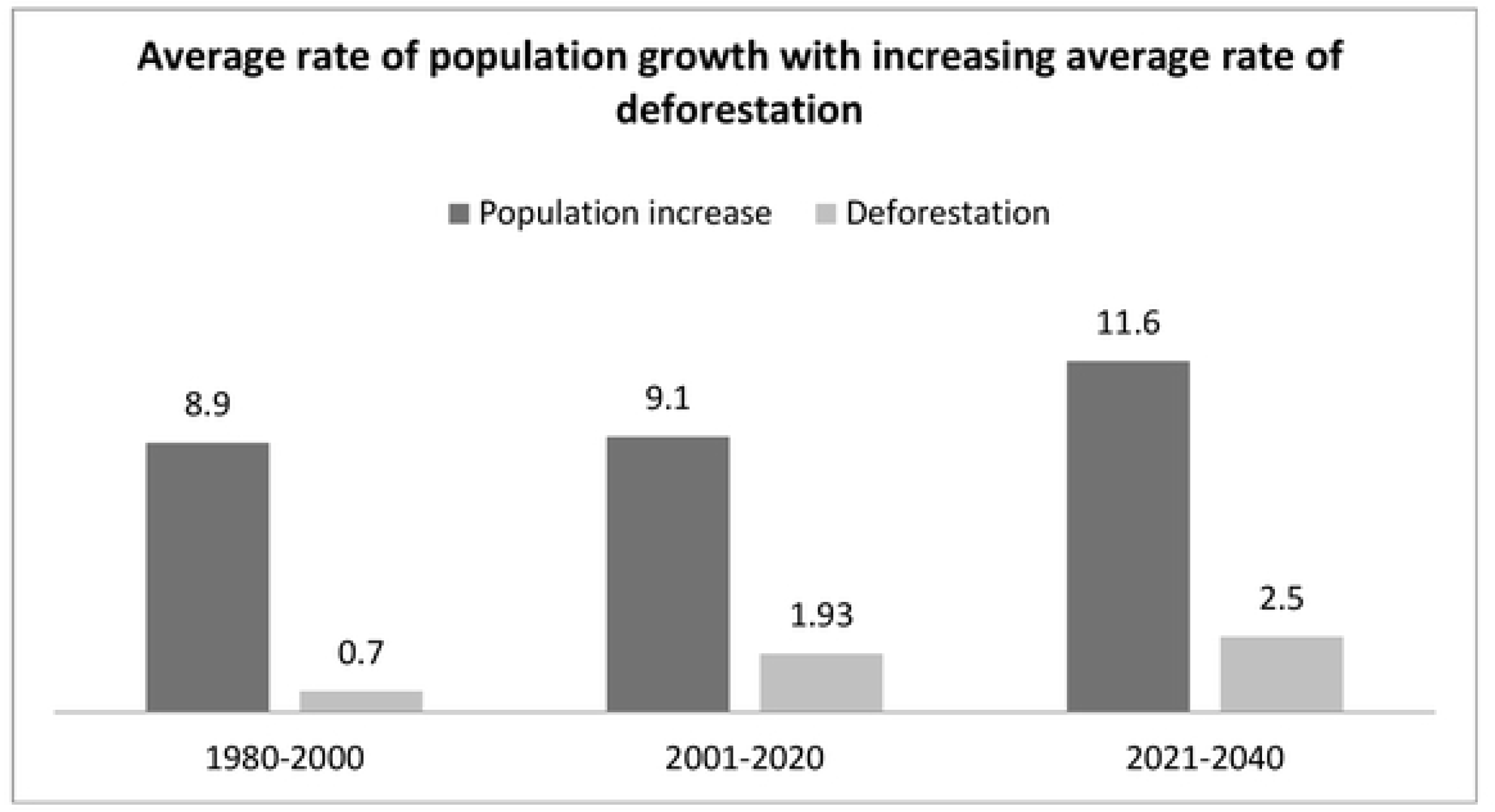
Average population growthand deforestation rates.

**Figure 5.**
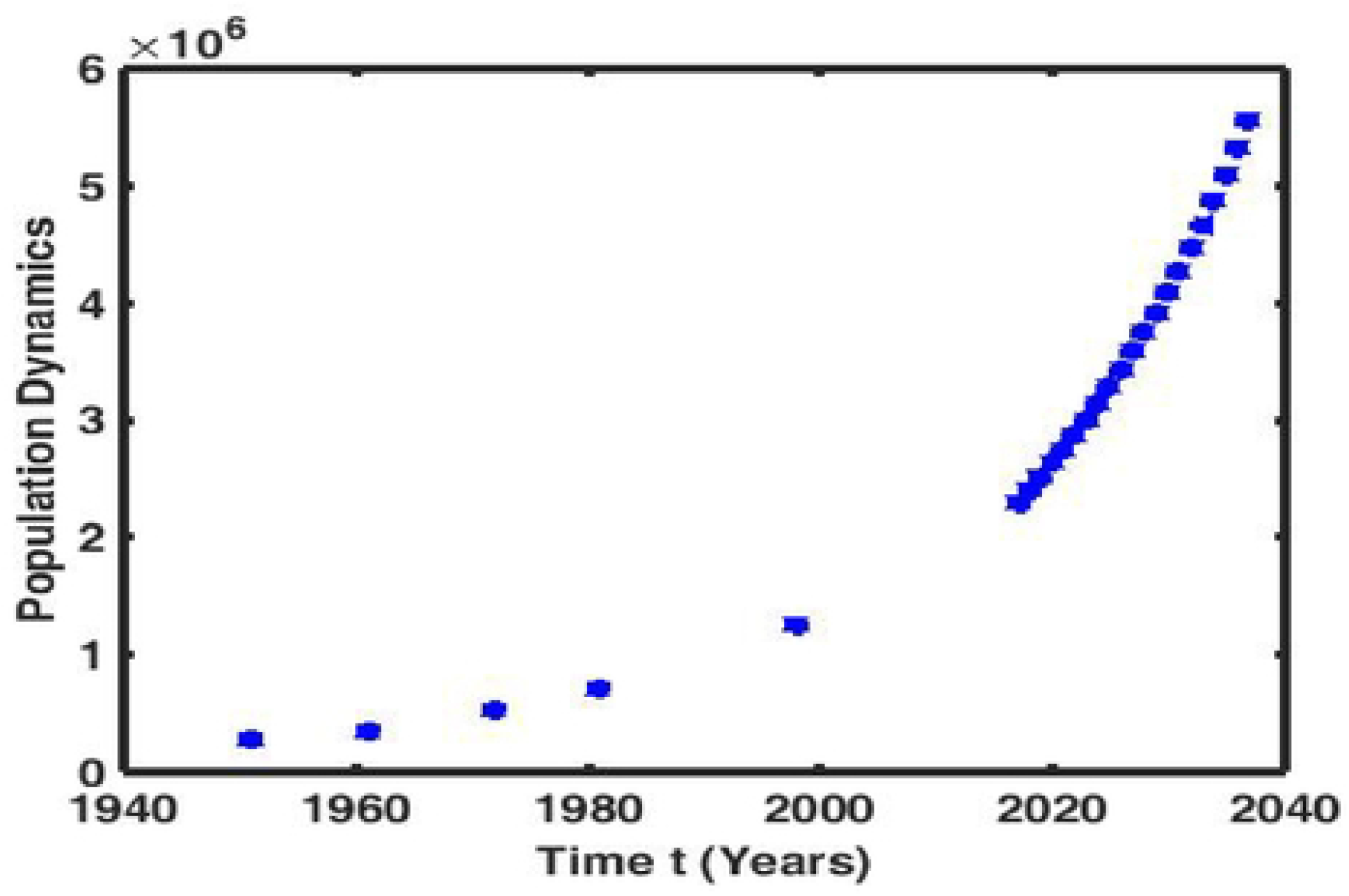
Decay model showing population increase in the year 2040.

**Figure 6.**
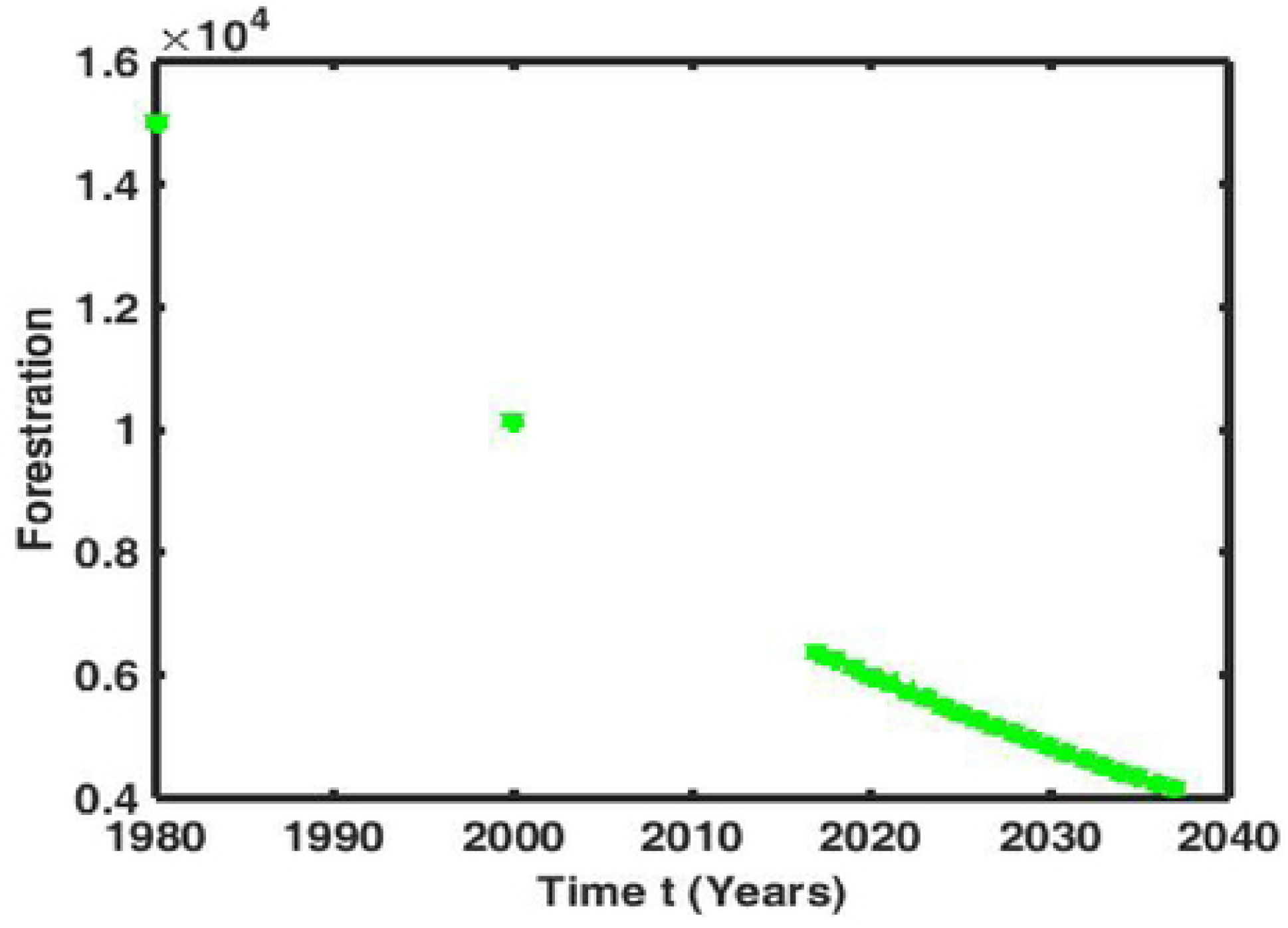
Decay model showing Forest Cover in 2040.

## 4. Discussion

Northern Pakistan’s Hindukush-Himalayan mountains are replete with temperate forests, and in addition to their many other benefits, these forests are among the country’s primary natural resources [38,39]. To meet their basic needs, local communities depend on forest resources, such as firewood, animal feed, building materials, food, and medicine. In addition, the sale of forest products like fuelwood, timber, medicinal plants, wild fruits, and vegetables provides a means of subsistence [14,27,40]. Despite the tremendous importance of forests for human society, forest ecosystems are threatened due to increasing human density and the unsustainable use of natural resources [14,23,24]. It is essential to determine the current status forest ecosystem and identify the significant drivers of deforestation for sustainable forest management [16]. The use of GIS and RS and proven to be the most convenient source of studying land use changes over a large area [18-20]. However, terrain and atmospheric effects degrade the quality of the images, which can be corrected with ERDAS software [28]. The Normalized Difference Vegetation Index (NDVI) was used to delineate the forest cover from RS satellite imagery across considerable time intervals [29], where the Decay model of exponential growth rate was able to determine the future projection and correlation of two parameters [35]. Our findings sought to describe the phenomenon of increased deforestation that has occurred in the study area over the last 40 years. From October 1980 to October 2000, the rate of deforestation was 0.7% per year, reducing the total forest area from 15,003 hectares to 12,859 hectares. From October 2000 to October 2020, the total forest cover decreased to 7,894 hectares at a rate of 1.93 percent annually. From October 1980 to October 2020, the average deforestation rate was 1.4%, indicating that the rate of deforestation is significantly higher than that reported by many authors in the nearby Hindukush-Himalayan mountains [41-43]. Deforestation is a major ecological issue in Northern Pakistan’s Hindukush-Himalayas Mountains. The main drivers of deforestation is population density, distance from the main town, and administration boundary [42]

Our findings indicate that population growth is the primary cause of deforestation in the study area. According to the 2017 district census report [32], the population of Swat district increased from 0.7 to 2.3 million over the last 40 years at an annual rate of 9% [44]. The decay model projection shows that the Swat district population will increase exponentially to 5.4M at a rate of 11.6% per year and predict a further loss of 2.5% per annum.[41] Population growth indicates that forest resources will be more at risk if long-term management planning in the study area is not implemented [45,46]. Numerous authors have previously reported on the severe effects of anthropogenic activities resulting from population growth. According to our findings, the population density of the study area will increase exponentially to 5.4 M with an average annual increase rate of 11.6% by 2040, while the forest density will further decrease to 4000 ha, as predicted by the decay model. The local people have limited sources of income, and non-timber forest products (NTFPs) are their primary income source. Continued practices of deforestation will further worsen their poverty and life standard. The carrying capacity of forests declines at a rate proportional to the pressures exerted on them by a growing population; nevertheless, an equilibrium level of forest cover can be maintained as a function of the population [47]. Unplanned and uncontrolled urbanization that has led to overpopulation in urban areas has had a catastrophic effect on the structure and function of ecosystems and has upset the natural ecological equilibrium [48]. Human population growth is associated with numerous anthropogenic activities that substantially contribute to deforestation, such as socioeconomic conditions that lead to illegal logging, agricultural expansion to feed hungry mouths, and increased fuel wood collection to meet rising energy demands [49]. Although ecotourism determines the economic value of the forest ecosystem (50), unsustainable tourism disturbance impacts the forest vegetation, and forest sites exposed to unsustainable tourism affect vegetation structure and diversity [51]. It is evident from different studies that unsustainable ecotourism have a negative impact on the environment including biodiversity and forest cover. Excessive visitors may disturb the fragile soils, vegetations and can cause wildlife conflicts [52].

Thus, it is essential to improve the conservation effort [42]. Priority areas for biodiversity conservation should be identified based on human population pressure, habitat status, and management activities [45]. The confluence of climate change and human population pressure threatens the resilience of ecosystem services [46]. Reforestation programs will be necessary to increase the amount of forest, and local communities should be urged to use forest resources sustainably. Alternative methods should also be introduced to lessen the demand for forest resources [4,52].

## 5. Conclusions

Deforestation threatens forest ecosystems worldwide, particularly in developing nations, causing a negative impact on the environment and the economy. According to local studies and international statistics, Pakistan has experienced significant deforestation, and the Malam Jabba forests of District Swat are no exception. To detect forest cover change over the past three decades, multispectral satellite images were analyzed using geospatial information systems (GIS) and remote sensing (RS) techniques to evaluate the effects of various anthropogenic activities in the study area. Supervised land cover classification and normalized difference vegetation indices (NDVIs) were applied to Landsat 5, 7, and 8 geospatial satellite images from 1980, 2000, and 2020. Our findings show that in the past two decades, the study area’s population has increased at a rate of 9.1% per year, which has led to an increase in anthropogenic activities such as road construction, hotel expansion, housing construction, fuel and timber wood extraction, overgrazing, poorly managed ecotourism, war and conflict, and shifting socioeconomic structure. Between 2000 and 2020, the annualized rate of deforestation increased from 0.7% to 1.93 % due to population growth-induced ecological imbalances. According to the exponential decay model, the District Swat’s population will increase exponentially to 5,7 million by 2040, at an annual rate of 11.6%. Increasing human activities will further impact the forest cover as the human population increases. According to the decay model, the forest cover in the study area will decrease to 4,000 hectares at a rate of 2.5 due to increasing population density and human activity. To ensure the long-term viability of natural forest resources, it is advised that local citizens, non-governmental organizations (NGOs), and the government departments responsible for conservation and management should work together to ensure community-based sustainable forest management. The government should provide alternative timber and fuel wood options to the local community. Awareness of the local community is essential for sustainable ecotourism, using natural resources, and successful joint forest management programs. The government should designate as a site for in-situ conservation the moist temperate forest of the study area, which contains timber trees, abundant floral diversity, and economically significant medicinal plants with sustainable ecotourism which will help to conserve these valuable forests.

## Author Contributions

“Conceptualization, Naveed Alam; writing—original draft preparation, methodology, software, validation, Zahid Ullah; formal analysis, investigation, Bilal Ahmad.; resources, data curation, Kashmala Syed writing—review and editing, Naveed Alam;, writing—original draft preparation, Ahmad Ali; visualization, Kashmala Syed.; supervision, project administration, Naveed Alam; funding acquisition, Naveed Alam. All authors have read and agreed to the published version of the manuscript.”

## Institutional Review Board Statement

Not applicable

## Data Availability Statement

Data are available upon reasonable request

## Acknowledgments

We are grateful to the Higher Education Commission (HEC) of Pakistan for funding this research project through Start-up Research Grant Program (SRGP) – R&D Division No: 21-1095/SRGP/R&D/HEC/2016

